# Synaptic vesicle characterization of iPSC-derived dopaminergic neurons provides insight into distinct secretory vesicle pools

**DOI:** 10.1101/2024.02.22.581435

**Authors:** Kenshiro Fujise, Martin Shaun Rosenfeld, Nisha Mohd Rafiq

## Abstract

The impairment of dopaminergic (DA) neurons plays a central role in the development of Parkinson’s disease. Evidence for distinct populations of synaptic vesicles (SVs) differing in neurotransmitter content (glutamate versus dopamine) has been attributed to differences in trafficking pathways and their exocytosis kinetics. However, the molecular and ultrastructural organization of the two types of vesicles remains poorly understood. Here we examined the development of axonal varicosities in human iPSC-derived DA neurons and glutamatergic neurons (i^3^Neurons). While i^3^Neurons are comprised of 40-50 nm small clear SVs, DA neurons are predominantly comprised of large pleiomorphic vesicles including empty and dense core vesicles, in addition to the classical SVs. The large vesicles were positive for VMAT2, the monoamine vesicular transporter responsible for loading dopamine, and are distinctly larger in size and spatially segregated from the VGLUT1/2-positive vesicles when expressed in an ectopic SV-like organelle reconstitution system. Moreover, these VMAT2-positive vesicles were also colocalized to known SV markers such as Rab3, SCAMP5, VAMP2, SV2C and can be clustered by the matrix protein synapsin. Our results show that DA neurons display inherent differences in their populations of neurotransmitter-containing secretory vesicles, and iPSC-derived neurons are powerful models for the study of presynaptic structures.

## INTRODUCTION

The selective degeneration or loss of dopaminergic (DA) neurons in the brain plays a central role in the development of Parkinson’s disease (PD). Several studies have directly linked abnormalities in synaptic vesicle (SV) recycling as critical contributors to the preferential vulnerability of DA neurons, particularly in the development of early-onset parkinsonism (EOP)(Cao et al., 2020; Cao et al., 2017; Ng et al., 2023; Vidyadhara et al., 2023). DA neurons are known for their large and highly arborized axon terminals comprising of dense varicosities innervating the striatum (Bolam and Pissadaki, 2012; Matsuda et al., 2009; Pacelli et al., 2015). Moreover, DA neurons have an innate ability to fire in two patterns: tonic (4-5 Hz) and bursting (∼20Hz) phases (Howe and Dombeck, 2016). Since the type of firing pattern can drastically affect neurotransmitter release (Koranda et al., 2014; Sulzer et al., 2016), understanding the spatial architecture of DA synapse is crucial in elucidating the preferential vulnerability of these neurons to synaptic dysfunction.

Synapses can be broadly categorized into classical or neuromodulatory depending on the type of neurotransmitters released. While classical chemical synapses for glutamate and γ-aminobutyric acid (GABA) are fast and target receptors in nanoscale proximity (presynaptic terminals and post-synaptic densities), neuromodulators such as dopamine operate at different spatiotemporal scales (Liu et al., 2021). In fact, DA neurons are thought to have predominantly in-passing presynaptic boutons with no clear postsynaptic component in the brain, and to a lesser extent the classical chemical synapses (Hattori et al., 1991; Tritsch et al., 2012; Wildenberg et al., 2021).

A crucial aspect that is lacking in the synaptic biology of DA neurons is the organelle(s) responsible for dopamine release. The storage of monoamines (serotonin, dopamine, histamine, adrenaline, and noradrenaline) is performed by the vesicular monoamine transporters (VMATs) 1 and 2. Both transporters exhibit tissue-specific abundance, with VMAT1 in neuroendocrine while VMAT2 found in DA and serotonin-containing neurons of the central nervous system (Deutch, 2013; Erickson et al., 1996). Unlike glutamate and GABA transmitters which are primarily stored in small clear synaptic vesicles (SSVs), the vesicles storing dopamine are reportedly heterogenous in their identities. For instance, VMAT2, the primary transporter for neuronal dopamine, is described to localize on SSVs and secretory dense core vesicles (DCVs) (Nirenberg et al., 1996; Nirenberg et al., 1995) or exclusively in DCVs (Melissa et al., 1997; Tao-Cheng et al., 1995). The presence of VMAT2 in secretory vesicles is consistent with the localization of VMAT1 on DCVs of the adrenal chromaffin cells (Kobayashi, 1977; Tao-Cheng et al., 1995; Yasothornsrikul et al., 2003).

However, little is known on the role of secretory vesicles on dopamine release in DA neurons. Moreover, evidence for SV transmitter pleiomorphism in DA neurons is primarily attributed to electrophysiological and behavioral analysis of experiments relating to transmitter co-release mechanisms (El Mestikawy et al., 2011; Hnasko et al., 2010; Onoa et al., 2010; Pereira et al., 2016; Silm et al., 2019). Recent evidences have also pointed to distinct variations in sorting pathways for VMAT2- and vesicular glutamate transporter (VGLUT)-containing vesicles, highlighting intrinsic differences in trafficking mechanisms for different SV transmitter pools (Asensio et al., 2010; Jain et al., 2023; Silm et al., 2019). Hence, a detailed characterization of the vesicles secreting dopamine becomes even more critical in our understanding of synaptic malfunction in neurodegenerative diseases such as Parkinson’s disease.

Towards this aim, our work seeks to gain insight into the anatomical and molecular organization of SV transmitter pools in DA neurons. Using a combination of cellular-based systems including glutamatergic iPSC-derived neurons (i^3^Neurons), iPSC-derived DA neurons and co-cultures of iPSC-derived DA neurons with their striatal target medium spiny neurons (MSNs), we found that DA neurons form distinct vesicle pools of varying sizes and nature: classical SSVs, large vesicles and DCVs. Using an ectopic SV-like organelle reconstitution system in fibroblasts, we show that VMAT2-positive vesicles can be clustered by synapsin and are sorted in heterogenous vesicular populations distinct from the classical VGLUT-containing vesicle clusters. In DA neurons, vesicles positive for VMAT2 do not resemble bona fide SSVs in terms of morphology, size and clustering properties. Our findings provide evidence that DA neurons use different classes of vesicles for the secretion of dopamine and of glutamate during synaptic transmission.

## RESULTS

### Characterization and maturation of iPSC-derived DA neurons

We used previously described DA differentiation protocol (Bressan et al., 2021; Kriks et al., 2011) to derive DA neurons from iPSCs (See Methods). Analysis of the cultures by immunofluorescence showed that 30 days from the induction of differentiation, 76.23% ± 7.8 (mean ± S.D.) of the cells were positive for the neuronal marker βIII-tubulin, and 89.21% ± 6.7 (mean ± S.D.) of the βIII-tubulin-positive neurons were positive for the DA neuron marker tyrosine hydroxylase (TH) (Figure 1A and 1B). Western blot analysis confirmed expression of TH, as well as expression of the plasma membrane DA transporter (DAT) in these cells (hence referred to as DA neurons), but not in neurogenin-2 induced iPSC-derived glutamatergic cortical-like neurons (i^3^Neurons, day 19) used as a control (Figure 1C).

**Figure 1:**
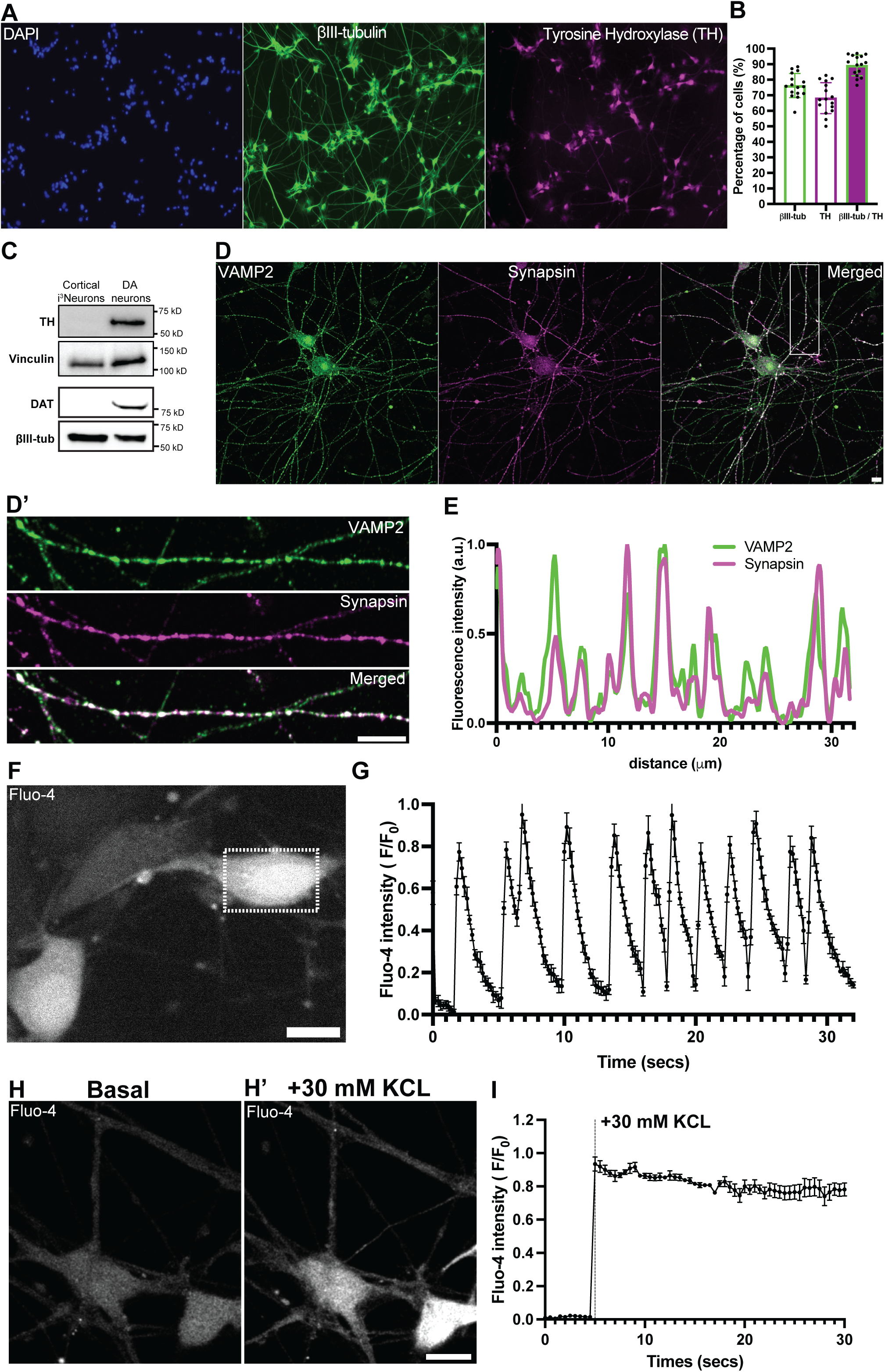
Characterization and maturation of iPSC-derived DA neurons. (A) Fluorescence images of iPSC-derived DA neurons (day 30) stained for DAPI to label nuclei (blue) and immunolabeled with antibodies directed against βIII-tubulin (green) and TH (magenta). Scale bar, 100 μm. (B) Percentage of cells positive for βIII-tubulin, TH or both, represented as mean ± SD pooled from five different experiments (n ≥ 50 cells per experiment). (C) Western blot of i^3^Neurons and iPSC-derived DA neurons immunoblotted for TH and DAT; vinculin and βIII-tubulin served as loading controls. (D) Representative fluorescence images of DA neurons (day 50) immunolabeled with antibodies against VAMP2 (green) and synapsin (magenta). (D’) High magnification of the boxed area in D is shown on the right. Scale bars, 10 μm. (E) Line scan intensity profile showing overlapping fluorescence signals of VAMP2 and synapsin in an axon of a DA neuron. (F) Live fluorescence imaging shows spontaneous calcium activity in DA neurons at day 50. Scale bar, 10 μm. (G) Graph depicting the corresponding fluorescence intensity across time from five different regions of the same neuron in the dotted boxed area in F represented as mean ± SD. (H, H’ and I) Live calcium imaging of DA neurons (day 50) (H) shows massive spike in calcium levels after high KCL stimulation (H’). Scale bar, 10 μm. (I) Graph showing the corresponding normalized Fluo-4 fluorescence intensities pre- and post KCL stimulation represented as mean ± SD.

Conversely, expression of typical SV markers, such as synaptophysin, synapsin, VAMP2 and Rab3, were not present at relevant level in DA neurons at 30 days, but robust expression was observed at 50 days by immunofluorescence, which revealed the expected punctate synaptic pattern (Figure 1D and 1E, Supplementary Figure 1A-C).

Characterization of synaptic function with the calcium indicator Fluo-4 showed basal spontaneous activity within the neuronal population (Figure 1F and 1G) while acute depolarization with high K+ stimulation (30 mM KCL) induced a synchronized rise in calcium levels in these neurons (Figure 1H and 1I). Collectively, these results indicate that iPSC-derived DA neurons exhibit synaptic properties typical of a neuron from day 50 onwards in culture.

### iPSC-derived DA neurons show vesicle pools distinct from those in i^3^Neurons

We next used transmission electron microscopy (EM) to examine the presence and type of secretory vesicles in neurite varicosities of cortical-like i^3^Neurons and DA neurons. In i^3^Neurons, numerous tightly-packed typical presynaptic clusters of ∼40 nm vesicles were already clearly visible at day 19 post-differentiation (Figure 2A-D). In DA neurons, the appearance of clusters of similar vesicles along axonal processes was much delayed relative to i^3^Neurons (Figure 3), consistent with the delayed expression of typical SV marker proteins (Figure 1 and Supplementary Figure 1). In favorable sections, many such clusters were clearly anchored to a region of the plasma membrane directly opposed to a region of neighboring cells positive for a thick or thin plasma membrane undercoating, confirming that they represent synaptic sites. Moreover, three morphologically distinct populations of such clusters were observed.

**Figure 2:**
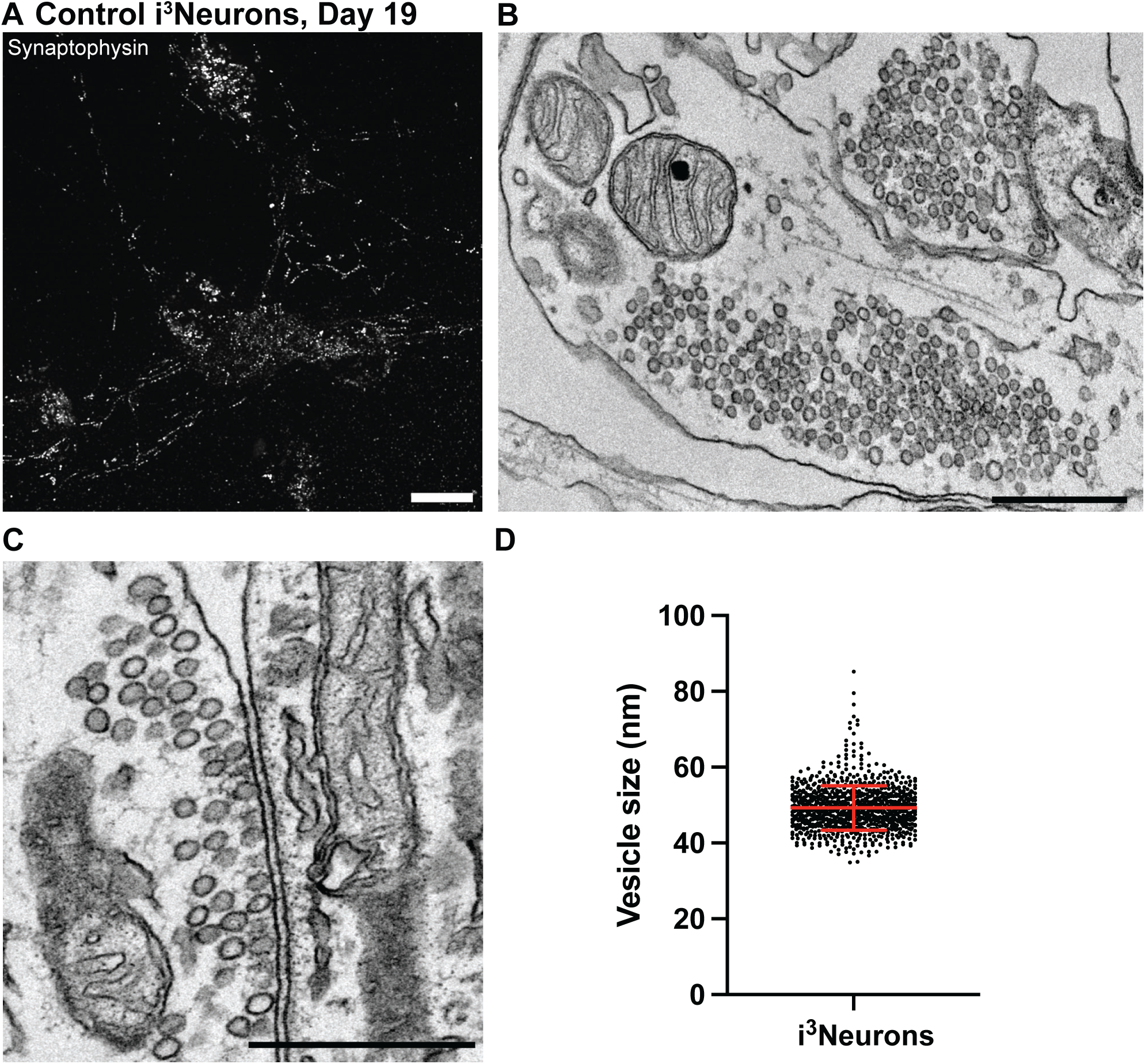
i^3^Neurons show presence of classical SSVs in axonal varicosities. (A) Fluorescence image of control i^3^Neurons (day 19) immunolabeled with antibody directed against synaptophysin. Scale bar, 10 μm. (B and C) Representative EM images showing numerous SSV clusters in presynaptic structures (B) of i^3^Neurons and the presence of electron dense postsynaptic densities in these cultures (C). Scale bars, 500 nm. (D) Quantification of SV size in i^3^Neurons represented as mean ± SD pooled from ≥ 878 synaptic vesicles.

**Figure 3:**
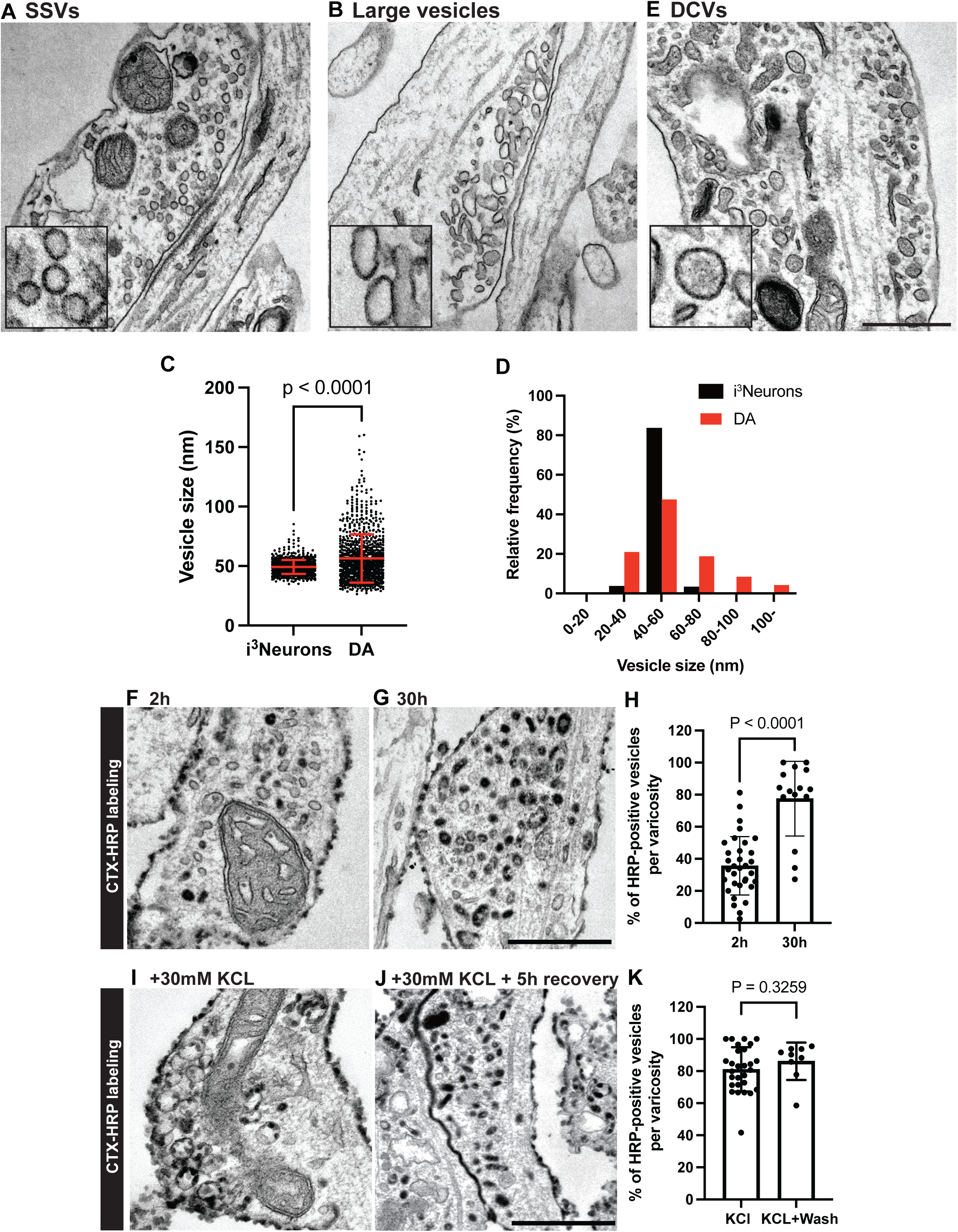
iPSC-derived DA neurons show size pleiomorphism in SV pools. (A-E) Representative EM images of DA neurons (day 50-55) differentiated from KOLF2.1 iPSCs show presence of SSVs (A), large vesicles (B) and DCVs (E) in classical synapses or bouton-like structures. Scale bar, 500 nm. Quantification of clear empty vesicles in i^3^Neurons (from figure 2D) and DA neurons are shown in dot plot (C) and frequency histogram (D). The dot plot is represented as mean ± SD pooled from ≥ 1723 vesicles. (F and G) EM images of DA neurons (day 50-55) incubated with CTX-HRP for 2 hours (F) and 30 hours (G) show dark staining of both small and large vesicles. Dark stainings indicate HRP-reactive labeling of vesicles originating from the plasma membrane. Scale bar, 500 nm. (H) Percentage of HRP-positive vesicles per varicosity represented as mean ± SD pooled from ≥448 vesicles in bouton-like structures. (I and J) High K+ stimulation induced stimulus-dependent bulk endocytosis, as a result most of these organelles were positive for HRP (I). During the recovery period (5 hours), bulk endosomes are converted to SSVs and large vesicles that were positive for HRP (J). Scale bar, 500 nm. (K) Percentage of HRP-positive vesicles per varicosity represented as mean ± SD pooled from ≥400 vesicles in bouton-like structures.

In one population, the ∼ 40 nm vesicles were round as in i^3^Neurons (Figure 3A, see also Figure 2B and C), and the post-synaptic membrane had the thick undercoating typical of post-synaptic densities of asymmetric synapses. These synapses are likely glutamatergic, consistent with the reported property of DA neurons *in situ* to form VGLUT-positive glutamatergic synapses (Silm et al., 2019; Zhang et al., 2015).

In another population, clusters of irregularly shaped vesicles with a clear content and around 60-100 nm in diameter were frequently observed (Figure 3B). These vesicles were tightly packed, completely segregated from other organelles and generally localized in outpocketing of neuronal processes, reminiscent of bouton-like structures lacking clear postsynaptic densities. Interestingly, appearance of these large vesicles has been observed in DA axonal boutons of the mouse nucleus accumbens (Wildenberg et al., 2021), whose identity has yet to be characterized. Analysis of the vesicle size distribution of both populations in DA neurons showed an average diameter of 54.9 ± 20.5 nm (mean ± S.D.), in comparison to those in i^3^Neurons (49.3 ± 5.8 nm and mean ± S.D., Figure 3C and D).

In addition to the two population of SV clusters described above, another vesicle population observed in DA neurons along cell processes was represented by 60-100 nm round, oval or irregularly shaped larger vesicles with an electron dense content. These vesicles were enriched in neurite varicosities but were much sparser than the other two populations described above. Their appearance suggests that they represent neuro-peptide containing large dense core vesicles (DCVs) (Figure 3E). The occurrence of DCVs was previously observed in nigrostriatal axons (Melissa et al., 1997). Their presence is of special interest, as in cells specialized for amine secretion, such as chromaffin cells of the adrenal gland, amines are stored in DCVs that also contain a variety of peptides.

To determine whether both small and large vesicles with clear contents are endocytic organelles, cholera toxin-horseradish peroxidase (CTX-HRP) was added to unstimulated, K+ stimulated, and during recovery from K+ stimulation. CTX-HRP labels endocytic organelles from the cell surface and is an efficient marker for tracing recently formed SVs. Incubation with CTX-HRP for 2 hours revealed a few vesicles positive for HRP most likely due to spontaneous neuronal activity, and a further increase in the number of HRP-positive vesicles after 30 hours (Figure 3F-H). Notably at 30 hours, most of the vesicles including the small and large ones were positive for HRP (Figure 3G). High K^+^ stimulation for 90 seconds resulted in massive formation of large HRP-positive vacuoles previously recognized as bulk endosomes (Figure 3I), often detected in the presynapse of mouse primary neuronal cultures after high K^+^ stimulation (Wu et al., 2014). Following recovery from high K^+^ stimulation, the majority of SVs were positive for HRP, indicating that the large pool of newly-formed SVs originated from the large vacuoles induced by high K^+^ stimulation (Figure 3J-I).

### Confirming the presence of large vesicles and DCVs in axon terminals

The presence of organelles reminiscent of large vesicles and DCVs which could be storage sites for DA raises the possibility that at least a pool of DA secreted from DA neurons in the striatum may be released from these organelles. As in neurons, these vesicles can be present both in dendrites and axons, we wanted to confirm that the neurites of DA neuron containing them were axons. To address this question, we used a microfluidic compartmentalization device in which neurons are seeded in one chamber but can extend axons to another chamber through long microchannels (640μm in length, Figure 4A, Supplementary Figure 2A and B).

**Figure 4:**
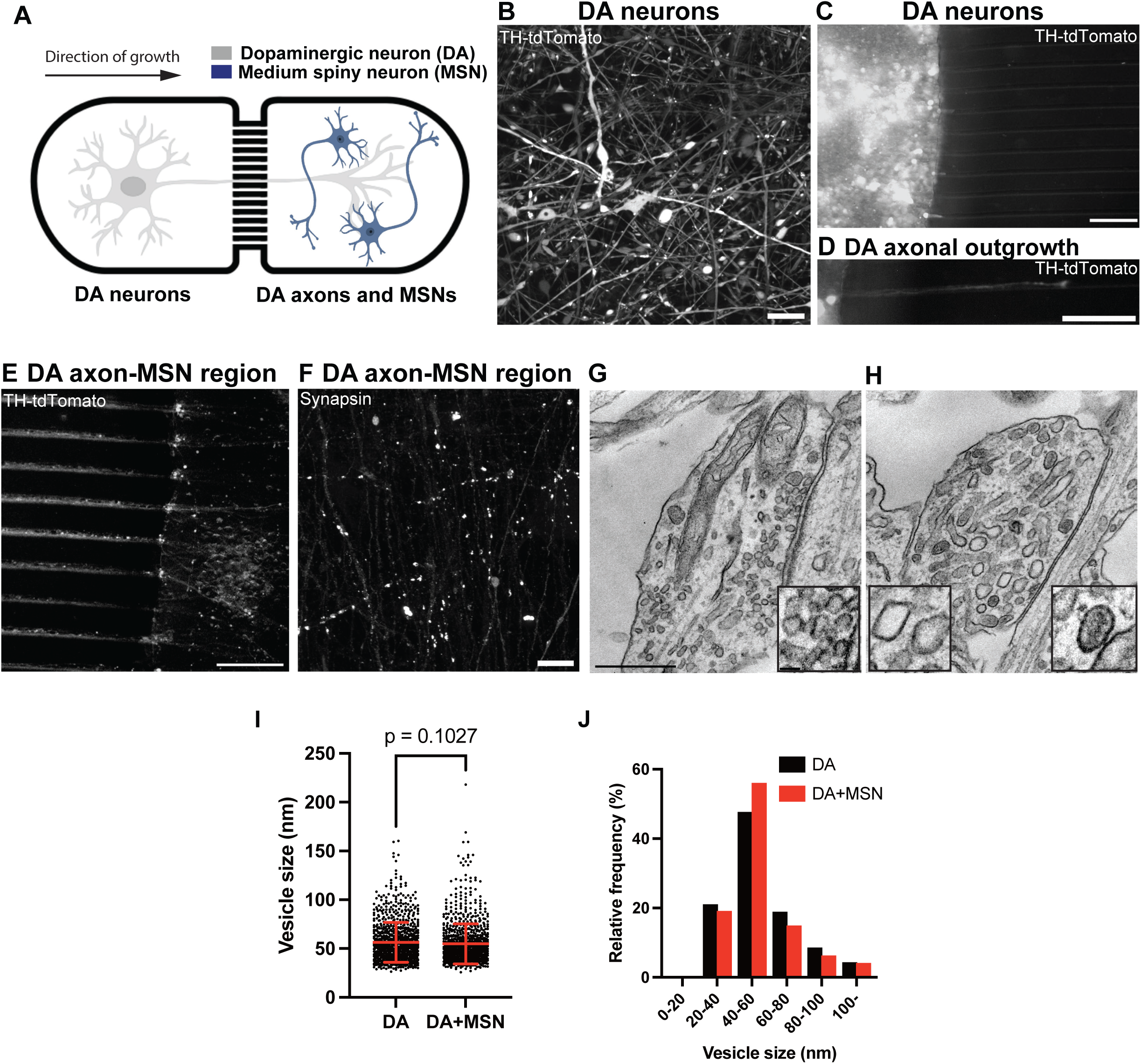
Co-culturing of iPSC-derived DA neurons with their striatal target neurons show size pleiomorphism in SV pools. (A) Diagram showing a schematic view of iPSC-derived DA neurons and iPSC-derived medium spiny neurons (MSNs) co-cultured in the microfluidic device. (B) Representative fluorescence image of endogenous tdTomato-tagged TH DA neurons (day 30) show robust TH labeling (white). (C and D) Fluorescence image of endogenous tdTomato-tagged TH DA neurons (white) plated on one side of the chamber (C), with a representative axon migrating through a microfluidic channel (D). Scale bars, 100 μm. (E) Fluorescence image of a representative area of the MSN-seeded chamber where only axons of the tdTomato-TH-positive DA neurons (white) have innervated the MSN-containing chamber (unlabeled). Scale bar, 100 μm. (F) Immunolabeling with antibody directed against synapsin show presence of synapses in this region (white), which could come from either DA-MSN or MSN-MSN contacts. Scale bar, 10 μm. (G and H) Representative EM images show presence of bona fide SSVs in a classical synapse (G) and large-sized vesicles (including clear and DCVs, see insets) in a presynaptic bouton-like structure (H). Scale bar, 500 nm. (I and J) Quantification of the size of these vesicles is shown in dot plot (I) and frequency histogram (J). The dot plot is represented as mean ± SD pooled from ≥ 1603 vesicles in bouton-like structures. Note that the vesicle pools in DA-MSN co-cultures were similar to those of single DA cultures only.

DA neurons detached from a 30-days old culture were plated in one of the chambers and allowed to grow their axons through the microchannels. To facilitate visualization of the axons, DA neurons derived from tdTomato-tagged TH iPSCs were used to track axonal outgrowth in the microchannels (Figure 4B-D). Moreover, to facilitate axonal outgrowth and formation of synapses, 10 days after plating the DA neurons, iPSC-derived striatal medium spiny neurons (MSNs), which we confirmed to be positive for DARPP32 immunoreactivity by western blotting and immunofluorescence (Supplementary Figure 2C-E), were seeded in the target chamber. A major synaptic target of DA neurons in the striatal region (nigrostriatal pathway) is the medium spiny neurons (MSNs), which represent 90% of the striatal neuronal population (Zhou, 2020). Following an additional 10 days, anti-synapsin immunofluorescence of the target chamber yielded abundant puncta revealing presence of axon varicosities (Figure 4E and F). Importantly, EM revealed the same abundant presence of the organelles reminiscent of irregularly shaped large vesicles and DCVs, revealing that these organelles populate axons (Figure 4G to J).

### Distinct localization of VMAT2 and VGLUT2 in DA neurons

VMAT2 is the vesicular transporter responsible for the loading of dopamine into secretory vesicles of nigrostriatal DA neurons. A non-overlapping localization of VMAT2 and VGLUT2 is in fact supported by previous studies of axons of DA neurons in the ventral striatum and primary cultures of DA neurons (Hnasko et al., 2010; Silm et al., 2019; Zhang et al., 2015).

To further validate this difference, we assessed the localization of VMAT2 and VGLUT2 in our iPSC-derived DA neurons at 50-55 days of differentiation, i.e. stage when synapses have matured (see Figure 1). Immunofluorescence of these cultures for VGLUT2 and synapsin demonstrated close colocalization of the two proteins in typical synaptic puncta (Figure 5A), confirming presence of glutamate-containing SVs. Since antibodies that yield reliable VMAT2 immunofluorescence were not available, expression of VMAT2 tagged at its C-terminus with either GFP or mCherry was used in this study. VMAT2-GFP colocalized with synapsin (like VGLUT2 vesicles), suggesting that they are indeed SVs and not just endosomes resulting from overexpression (Figure 5B). When DA neurons expressing tagged-VMAT2 were stained for VGLUT2, there was indeed only a partial overlap between VGLUT2 and VMAT2 signal (Figure 5C and D). In addition, we further validated the presence of endogenous VMAT2 and VGLUT2 in these neurons by western blot from day 30 and day 60 cultures (Figure 5E). Collectively, our results indicate that the types of transporters do not have the same localization, phenocopying what had been observed in primary neuronal cultures and striatal brain slices (Onoa et al., 2010; Silm et al., 2019).

**Figure 5:**
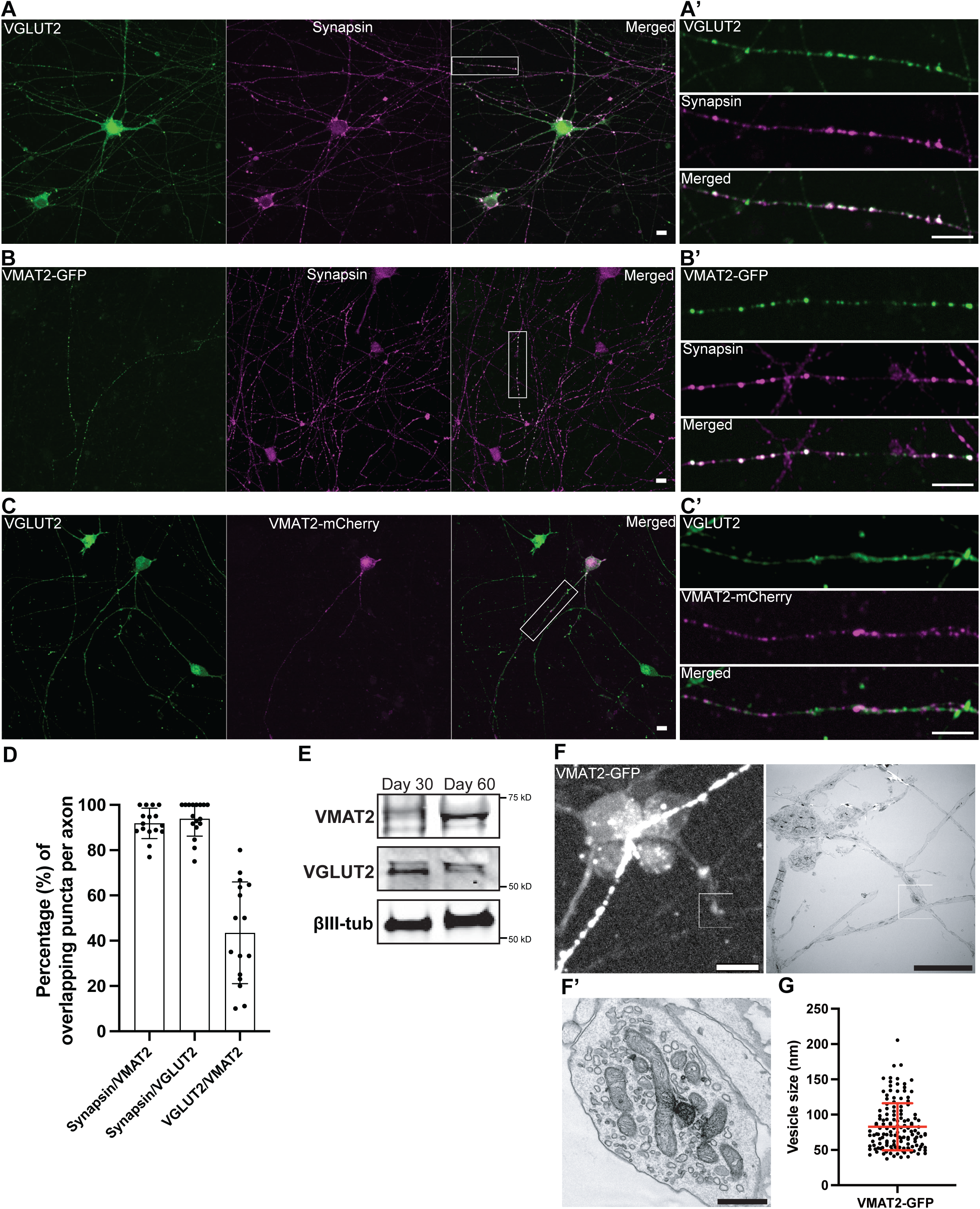
Fluorescent-tagged VMAT2 show minimal overlap with VGLUT2 in iPSC-derived DA neurons. (A) Fluorescence image of DA neurons (day 55) immunolabeled with antibodies directed against VGLUT2 (green) and synapsin (magenta). (A’) High magnification of an axon from the boxed area in A show overlapping signals (white) of VGLUT2 and synapsin. (B) Overexpression of VMAT2-GFP (green) in DA neurons (day 55) show overlapping fluorescence intensities with endogenous synapsin by antibody staining (magenta). (B’) High magnification of an axon from the boxed area is shown in B’. (C) Overexpression of VMAT2-GFP (green) in DA neurons (day 55) immunolabeled with antibody against VGLUT2 (magenta) show little overlap between these transporters. (C’) High magnification of an axon from the boxed area is shown in C. Scale bars, 10 μm. (D) Quantification of the overlapping signal punctas between synapsin/VMAT2, synapsin/VGLUT2 and VGLUT2/VMAT2 represented as mean ± SD, pooled from two independent experiments (n ≥ 5 cells per experiment). (E) Anti-VMAT2, anti-VGLUT2 and anti-βIII-tubulin (loading control) western blot of DA neurons at days 30 and 60. (F) Fluorescence image of DA neurons transfected with VMAT2-GFP (day 53, white) and the corresponding EM image (right) is shown on the right. Scale bars, 10 μm. (F’) High magnification of boxed areas show presence of clear large vesicles and the occasional presence of DCVs that were positive for VMAT2-GFP signals. Scale bar, 500 nm. (G) Quantification of VMAT2-GFP positive vesicles pooled from four EM images represented as mean ± SD.

To further understand the type of vesicles VMAT2 is localized on in these neurons, we first performed corelative light and electron microscopy (CLEM) for VMAT2 in DA neurons.

Towards this aim, VMAT2-GFP was transfected into DA neurons at day 50 on gridded glass dishes and visualized for VMAT2 fluorescence at day 53. VMAT2-GFP-positive structures were then identified on the gridded dish, and selected regions were processed for EM analysis (Figure 5F). EM observation revealed that clusters of VMAT2-GFP fluorescence in control DA neurons corresponded to a mixture of predominantly larger vesicles (>60 nm, Figure 5F’ and G), which are mostly empty and occasionally electron-dense, suggesting that VMAT2 proteins are indeed present on larger-sized secretory vesicles.

### Differential sorting of VMAT2 and synaptophysin when expressed in an ectopic system

To further assess differences in the intracellular sorting of VMAT2 relative to other SV proteins, we capitalized on a recent demonstration that clusters of SV-like organelles – condensates that appear as large droplets in fluorescence microscopy - can be generated in fibroblastic cells (COS7 cells) by exogenous expression of synapsin and synaptophysin (Park et al., 2021; Park et al., 2023). We had found that when additional bona fide SV proteins, such as VAMP2, SCAMP5, synaptotagmin 1, VGLUT1, VGAT1 and Rab3A are expressed together with synaptophysin and synapsin in these cells, they co-assemble with synaptophysin in the same vesicles. Thus, we examined whether also VMAT2, like these other proteins, can assemble into vesicles generated by synaptophysin expression.

First, we expressed VMAT2-GFP alone in COS7 cells and found that it localizes to Golgi complex and scattered vesicular puncta throughout the cytoplasm (Supplementary Figure 3A). When co-expressed together with mCherry-synapsin, VMAT2 co-assembled with synapsin into droplet-like condensates (Figure 6A and B) reminiscent of those generated by synaptophysin and synapsin (Figure 6C and D). However these condensates were composed of larger and irregularly shaped vesicles (81.1 ± 34.9 nm; mean ± S.D.), clearly different from those found in the synaptophysin-synapsin condensates (43.3 ± 8.2 nm ; mean ± S.D.) (Figure 6B, D, E and F). Strikingly, when VMAT2-GFP, synaptophysin and mCherry-synapsin were co-expressed together, VMAT2 vesicles and synaptophysin vesicles segregated from each other and assembled into distinct phases within the mCherry-synapsin phase, with synapsin being more concentrated (based on a higher fluorescence intensity) in the synaptophysin subphase (Figure 6G and 6H). Most interestingly, CLEM of cells co-expressing synaptophysin, synapsin and VMAT2 revealed that the two phases detectable by fluorescence correlated with two classes of vesicles: small SV-like vesicles in the synaptophysin phase and larger vesicles in the VMAT2 phase (Figure 6I). Moreover, either VGLUT2 (VGLUT2-GFP, Supplementary Figure 3B and C) or VGLUT1 (VGLUT1-GFP, Supplementary Figure 3D and E), when co-expressed with VMAT2 (VMAT2-FLAG), synaptophysin (untagged) and synapsin (mCherry-synapsin) showed selective preference of the glutamate transporters for the synaptophysin-synapsin condensates over the VMAT2-synapsin clusters. In addition, other SV proteins, such as VAMP2, Rab3, SCAMP5, synatotagmin-1 or SV2C (an SV protein highly expressed by nigrostriatal DA neurons), were all positively colocalized with the VMAT2-synapsin clusters (Figure 7A-E). Furthermore, while knockdown (KD) of AP3, an adaptor protein known to be important for the intracellular sorting of VMAT2 (Silm et al., 2019), prevented the formation of VMAT2-synapsin condensates (Supplementary Figure 4A-E) but did not impact the formation of synaptophysin-synapsin condensates (Supplementary Figure 4F and G), pointing to different sorting pathways for the two proteins. The separation of VMAT2-positive vesicles from vesicles that share greater similarity to bona fide SVs is consistent with the differential localization of VGLUT2 and VMAT2-postive vesicles in mouse DA neurons of the ventral striatum (Onoa et al., 2010; Silm et al., 2019; Zhang et al., 2015) and in iPSC-derived DA neurons as shown above. Note that while DA neurons have both large vesicles and DCVs, lack of a dense core in these vesicles is not unexpected, as fibroblastic cells do not express the molecular machinery to generate the peptide containing secretory granules of the classical regulated secretory pathway (De Camilli and Jahn, 1990). Hence it remains possible that VMAT2 is localized to both large clear and DCVs in DA neurons.

**Figure 6:**
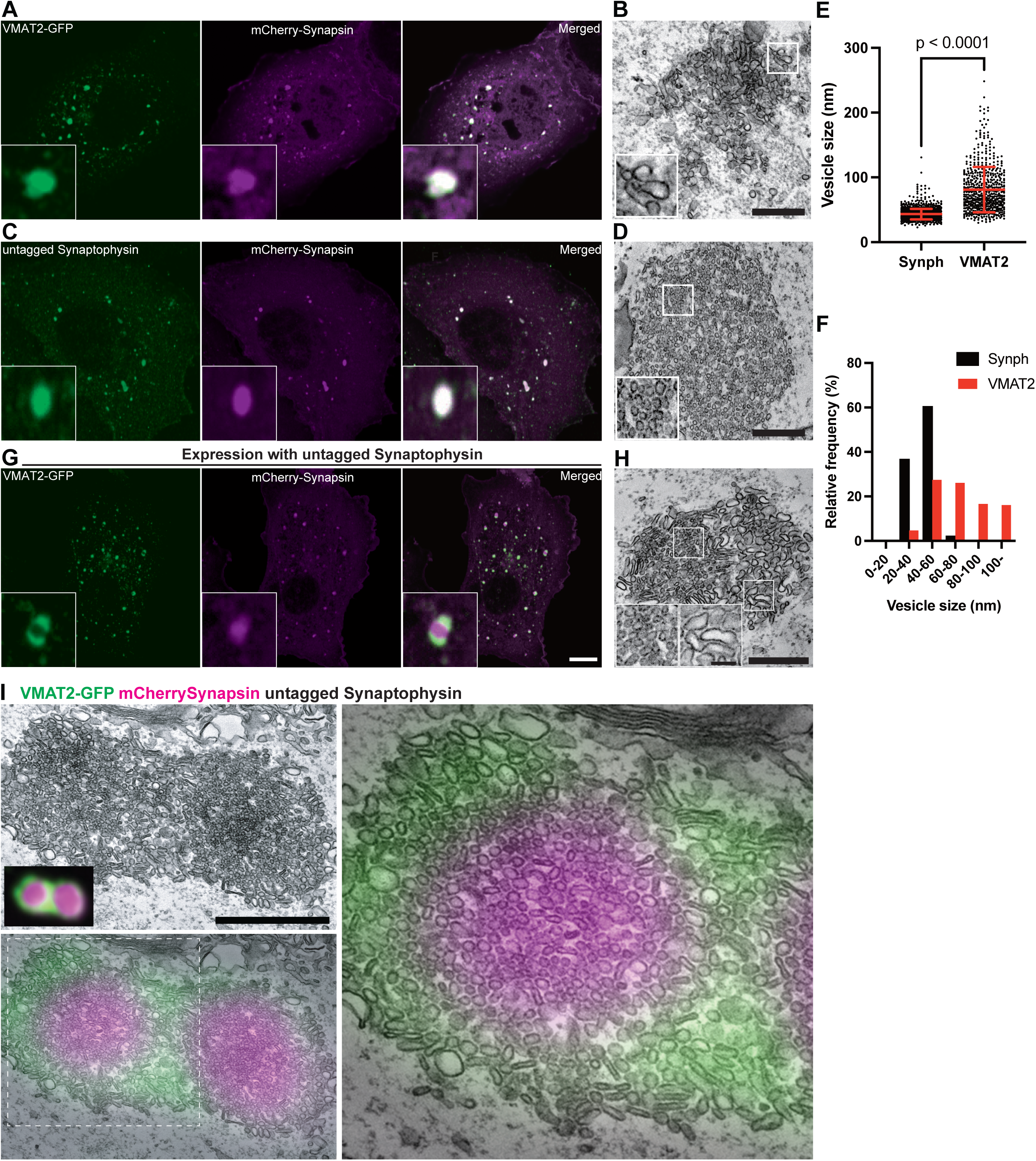
Co-expression of VMAT2-GFP with mCherry-synapsin results in clusters of large vesicles distinct from the synaptophysin-synapsin vesicle clusters. (A-D) Representative fluorescence and EM images of COS7 cells transfected with (A) VMAT2-GFP (green) and mCherry-synapsin (magenta) or (B) untagged synaptophysin (green) and mCherry-synapsin (magenta). Synaptophysin was visualized by antibody directed against it. Scale bars, 10 μm. The representative EM images (B and D) are depicted on the right of each panel. Scale bars, 500 nm. (E and F) Quantifications of individual vesicle size from the VMAT2-synapsin and synaptophysin-synapsin vesicle clusters are shown in dot plot (E) and frequency histogram (F). The dot plot is represented as mean ± SD pooled from ≥620 vesicles in vesicles clusters of four different cells. (G and H) COS7 cell triple co-transfected with synaptophysin (untagged), VMAT2-GFP (green) and mCherry-synapsin (magenta) show distinct clusters of vesicle pools (G). The representative EM image on the right (H) show variations in individual vesicle size between the two cluster populations. (I) Co-expression of synaptophysin, VMAT2-GFP (green) and mCherry-synapsin (magenta) in COS7 cells were fixed for correlative light-electron microscopy (CLEM). The synapsin phases include two distinct vesicle clusters containing synaptophysin (unlabeled but has strong synapsin labeling: magenta) and VMAT2-GFP (green) with lower synapsin fluorescence, which are comprised of small and large pleiomorphic vesicles, respectively.

**Figure 7:**
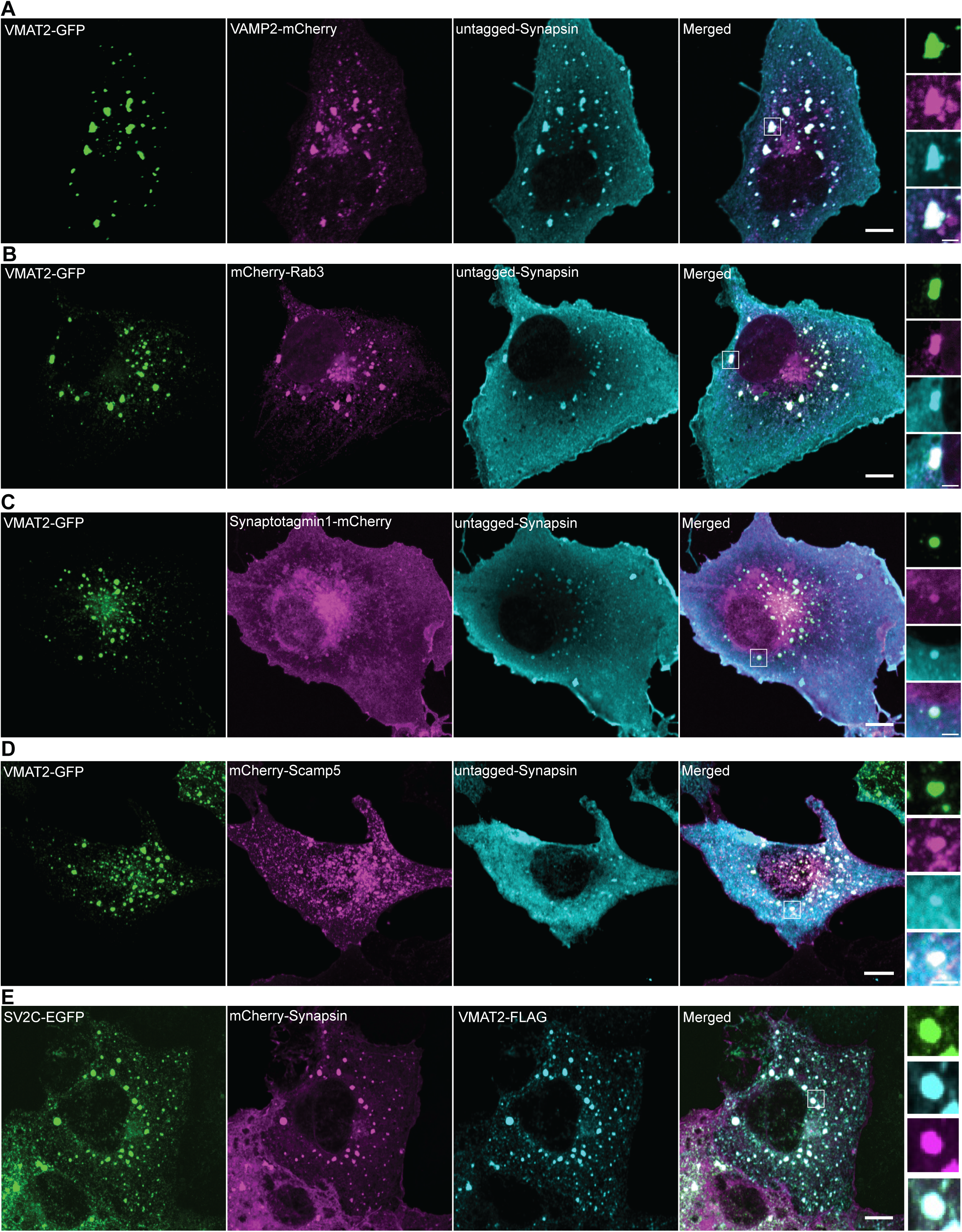
VMAT2 and synapsin vesicle clusters recruit other SV proteins. (A-E) COS7 cells were triple-transfected with untagged synapsin (cyan) and VMAT2-GFP (green) or mCherry-synapsin 1 (magenta) and VMAT2-Flag (cyan) with one of the following proteins as indicated: VAMP2-mCherry (magenta) (A), mcherry-Rab3A (magenta) (B), synaptotagmin 1-mCherry (magenta) (C), mCherry-SCAMP5 (magenta) (D), and SV2C-GFP (green) (E). Untagged synapsin and VMAT2-Flag were revealed by immunofluorescence. High magnifications of box areas are shown on the right for each panel. Scale bars, 10 μm; insets: 2.5 μm.

## DISCUSSION

Our results show a striking morphological difference in the SV pool populations in iPSC-derived DA neurons. In mice, DA nerve terminals were previously described to contain SVs positive for both VMAT2 and VGLUT2, necessary for the storage of dopamine and glutamate transmitters, respectively (Fortin et al., 2019; Hnasko et al., 2010; Onoa et al., 2010; Silm et al., 2019; Zhang et al., 2015). While both VMAT1 and VMAT2 are known to localize to secretory vesicles like DCVs in neuroendocrine cells (Li et al., 2005; Tao-Cheng et al., 1995; Weihe et al., 1994), it is generally believed that dopamine-containing vesicles in neurons are morphologically identical to the VGLUT- and VGAT-containing SV counterparts: clear vesicle clusters with a vesicular diameter of about 40 nm (Cao et al., 2017; Takamori et al., 2006).

We now demonstrate that DA nerve terminals show three pools of secretory vesicles: SSVs, large empty vesicles and DCVs in iPSC-derived DA neurons, while the same iPSCs differentiated into glutamatergic i^3^Neurons were comprised of predominantly SSV clusters. The large vesicles were positive for VMAT2 in DA neurons and in an SV-like cluster reconstitution system in fibroblasts. This finding opens the possibility of using iPSC-derived DA neurons as a synaptic model to gain new insight into the still open question of how DA neurons regulate neurotransmitter release in nerve terminals, which are predominantly comprised of non-classical synapses (Descarries et al., 1996; Hattori et al., 1991; Liu et al., 2021; Wildenberg et al., 2021).

An important result of our study, which was also previously documented in mouse and rat striata (Silm et al., 2019; Zhang et al., 2015), is the partial segregation of VGLUT2 and VMAT2-positive SVs in DA neurons. We show that VMAT2 localize on vesicles larger than bona fide SVs in iPSC-derived DA neurons and are also positive for synapsin. In addition, while vesicular transporters such as VGLUT1 and VGAT self-organize into small SV-like vesicle clusters by co-expressing synaptophysin and synapsin in fibroblasts (Park et al., 2021), the co-expression of VMAT2 with synaptophysin and synapsin resulted in the segregation of two distinct clusters: synaptophysin-synapsin and VMAT2-synapsin vesicle pools. The larger-sized VMAT2-positive vesicles in fibroblasts were strikingly reminiscent of those large vesicles (including DCVs) in DA neurons, which we further confirmed to be VMAT2-positive by CLEM. More importantly, co-expression of several SV proteins including Rab3, VAMP2, SCAMP5 and SV2C but not VGLUT1 or VGLUT2 were found to localize to the VMAT2-positive vesicles in fibroblasts, suggesting that both transporters are differentially sorted in cells. Furthermore, it has been previously reported that the lack of AP3 abolished the formation of VMAT2-postive vesicles but not the VGLUT2 vesicles in DA neurons (Silm et al., 2019), suggesting differential recycling pathways for both dopamine and glutamate transporters. Indeed, we showed that knockdown of AP3 in fibroblasts abolished formation of VMAT2-synapsin vesicle clusters, but not the synaptophysin-synapsin clusters.

More recently, large scale tracing of DA neurons in mouse nucleus accumbens using volume-based serial EM revealed presence of small and large vesicle clusters in DA axonal varicosities, predominantly lacking classical synapses (Wildenberg et al., 2021). These large vesicles were either clear or electron-dense (reminiscing those of DCVs) in nature. It is very likely that these vesicles, which were identified predominantly in non-classical synapses lacking postsynaptic densities, are VMAT2-positive, based on our findings in both iPSC-derived DA neurons and SV-like overexpression system in fibroblasts.

Collectively, our study demonstrates that, both in dopaminergic nerve terminals and in an exogenous SV-like organelle reconstitution system, major SV proteins and of VMAT2 are differentially sorted and represent an important step towards the elucidation of how dopamine transmitters are stored and released during synaptic transmission. iPSC-derived DA neurons would serve as a fantastic model to study how classical and non-classical synapses are interconnected to each other, a facet of dopaminergic neurobiology that is still insufficiently characterized.

## ACKNOWLEDGEMENTS

This work was supported in part by grants from the National Institutes of Health (NS036251), and the Parkinson’s Foundation (PF-RCE-1946) to Pietro De Camilli. N.M. Rafiq acknowledges support from the Deutsche Forschungsgemeinschaft (DFG, German Research Foundation, project number 335549539—GRK 2381) and core funding at the University of Tübingen (Interfaculty Institute of Biochemistry). For the purpose of open access, the author has applied a CC public copyright license to all Author Accepted Manuscripts arising from this submission.

We thank Peng Xu (Yale University, USA) for useful discussions and constructive criticism. Author contributions: N.M. Rafiq and K. Fujise conceptualized the project and wrote the manuscript. K. Fujise and N.M. Rafiq equally designed and performed all experiments. M.S. Rosenfeld provided technical assistance. All authors contributed intellectually to the project.

## MATERIALS AND METHOD

### Antibodies and plasmids

All VMAT2 and VGLUT1/2 constructs used in this study utilized a human codon-optimized sequence and were cloned using the Gateway recombination cloning approach. The following donor constructs were obtained from RESOLUTE consortium SLC collection on Addgene: pDONR221_SLC18A2 (#131997), pDONR221_SLC17A7(#131960), pDONR221_SLC17A6 (#131948), pDONR221-SLC6A3_STOP (#161326), pDONR221_SV2C (#132297). Briefly, VMAT2 (SLC18A2), VGLUT1 (SLC17A7), VGLUT2 (SLC17A6) and SV2C were tagged at the c-terminus with EGFP, mCherry or 2xFLAG in the following destination vectors pDEST-eGFP-N1(#31796), pDEST-mCherry-N1 (#31907) and 2xFlag-pDEST-C (#118372), respectively. The SLC6A3 (DAT) was n-terminally tagged with mCherry in the pDEST-CMV-N-mCherry (#123215) destination vector.

The following plasmids: untagged synaptophysin, untagged synapsin 1a, mCherry-synapsin 1a, VAMP2-mCherry, mCherry-Rab3a, mCherry-SCAMP5, Synaptotagmin1-mCherry were previously generated in the laboratory of Pietro De Camilli. Each construct was validated by DNA sequencing. All antibodies used in this study are listed in Table S1.

### Human iPSC culture, i^3^Neuron and DA differentiation

The following iPSC lines were obtained from the iPSC Neurodegeneration Initiative (iNDI) consortium and genome-edited by Jackson Laboratories (JAX): KOLF2.1, KOLF2.1 with the NGN2 cassette at the AAVS locus (used for the i^3^Neurons experiments) and tdTomato-tagged TH KOLF2.1 iPSCs. For the maintenance of iPSCs in culture, iPSCs were cultured on Geltrex (Life Technologies) coated dishes and maintained in Essential 8 Flex media (Thermo Fisher Scientific). The Rho-kinase (ROCK) inhibitor Y-27632 (EMD Millipore, 10 μM) was added to Essential 8 Flex media on the first day of plating and replaced with fresh media without ROCK inhibitor on the following day.

For i^3^neuronal differentiation, iPSCs were differentiated into cortical-like i^3^Neurons according to a previously described protocol based on the doxycycline inducible expression of Ngn2 (Fernandopulle et al., 2018). Briefly, iPSCs were dissociated with Accutase (Thermo Fisher Scientific) and re-plated at a density between 1.5–3 × 10^5^ cells on geltrex-coated dishes in induction medium [(KnockOut DMEM/F-12 (Thermo Fisher Scientific) containing 1% N2-supplement (Thermo Fisher Scientific), 1% MEM non-essential amino acids (Thermo Fisher Scientific), 1% GlutaMAX (Thermo Fisher Scientific) and 4 μg/mL doxycycline (Sigma-Aldrich)]. After 3 days, pre-differentiated i^3^Neurons were dispersed using Accutase and plated on 0.1 mg/ml poly-L-ornithine (Sigma-Aldrich) in borate buffer and 10 μg/ml laminin (Thermo Fisher Scientific) coated 35 mm glass-bottom dishes (MatTek) or 6-well plates (Corning) for imaging and immunoblotting, respectively. These i^3^Neurons were cultured and maintained in cortical medium (induction medium supplemented with 2% B27 (Thermo Fisher Scientific), 10 ng/mL BDNF (PeproTech), 10 ng/mL NT-3 (PeproTech) and 10 μg/mL laminin). Fresh cortical media was added to the existing media every 5 days. The iPSCs and i^3^Neurons were kept at 37°C with 5% CO2 in an enclosed incubator. A detailed protocol can be found at https://www.protocols.io/view/culturing-i3neurons-basic-protocol-6-n92ld3kbng5b/v1.

For the differentiation of iPSCs to DA neurons, we used the following protocols described in Kriks et al. (2011) and Bressan et al. (2021). Briefly, iPSCs were dissociated with Accutase (Thermo Fisher Scientific) and re-plated at a density of 8 × 10^5^ cells per well (of a 6-well plate) on geltrex-coated dishes in Essential 8 Flex media with Rock inhibitor. On the next day (Day 0 of differentiation), the media was replaced with knockout serum replacement (KSR) media containing 500nM LDN193189 (STEMCELL Technologies) and 10 μM SB431542 (STEMCELL Technologies). KSR medium is comprised of Knockout DMEM/F12 medium, 15% Knockout serum replacement (Thermo Fisher Scientific), 1% MEM NEAA, 1% glutaMAX, 0.1% 2-mercaptoethanol (Thermo Fisher Scientific) and 0.2% penicillin-streptomycin (Thermo Fisher Scientific). Starting the following day (Day 1) 75% of the differentiation medium was replaced with new medium each day from day 1 to day 15, then every 2 days until day 20. For days 1-4, KSR medium containing 500 nM LDN193189, 10 μM SB431542, 200 ng/ml SHH C25II (R&D Systems), 2 μM Purmorphamine (Cayman Chemical Company) and 100 ng/ml FGF-8b (PeproTech) was added daily, supplemented by the addition of 4 μM CHIR99021 on day 3 and 4. For days 5 and 6, a mixture of 75% KSR + 25% N2 medium also containing 500 nM LDN193189, 10 μM SB431542, 200 ng/ml SHH C25II (R&D Systems), 2 μM Purmorphamine (Cayman Chemical Company), 100 ng/ml FGF-8b (PeproTech) and 4 μM CHIR99021 (Tocris) was added to the cells followed by equal amounts of KSR and N2 media on days 7-8, and 25% KSR + 75% N2 media on days 9-10 also containing 500 nM LDN193189, 10 μM SB431542, 200 ng/ml SHH C25II and 4 μM CHIR99021. The N2 medium is comprised of Neurobasal Plus media (Thermo Fisher Scientific), 2% B27 supplement without vitamin A (Thermo Fisher Scientific), 1% N2 supplement, 1% glutaMAX and 0.2% penicillin-streptomycin. For days 11-20, complete NB/B27 medium was added to cells, with the addition of 4 μM CHIR99021 on days 11 and 12 only. Complete NB/B27 medium is comprised of N2 medium (without the N2 supplement) and the following components: 20 ng/ml BDNF (PeproTech), 0.2 mM ascorbic acid (Sigma-Aldrich), 20 ng/ml GDNF (PeproTech), 0.5 mM db-cAMP (Sigma-Aldrich), 1 ng/ml TGFβ3 (R&D Systems) and 10 μM DAPT (Cayman Chemical Company). After 20 days of culture, DA progenitor cells were frozen in Synth-a-freeze cryopreservation media (Thermo Fisher Scientific) and stored at -80^0^C or liquid nitrogen.

For long-term culture of DA neurons, cells were re-plated on 0.1 mg/ml poly-L-ornithine in PBS (Sigma-Aldrich) and 10 μg/ml laminin (Thermo Fisher Scientific) coated 35 mm glass-bottom dishes (MatTek) or 6-well plates (Corning) for imaging and immunoblotting, respectively. These neurons were cultured and maintained in complete NB/B27 medium followed by the addition of 0.1% anti-mitotic inhibitor (Supplement K, Brainxell) at day 25 to terminate division of non-neuronal cells. Fresh NB/B27 medium was added to the existing plates or dishes every 7 days and kept at 37°C with 5% CO2 in an enclosed incubator.

### Cell culture and transfections

COS7 cells were grown in DMEM (Thermo Fisher Scientific) supplemented with 10% FBS (Thermo Fisher Scientific) and 1% penicillin-streptomycin. Cells were kept at 37°C with 5% CO2 in an enclosed incubator. For transfection of COS7 cells, 1 μl Lipofectamine™ 2000 Transfection Reagent (Invitrogen) was used with the respective plasmids and visualized within 24-48 hours. For both i^3^Neuron and DA neuron transfections, plasmids were transfected with 4 μl of Lipofectamine™ Stem Transfection Reagent (Invitrogen) and visualized at least 48 hours later. For siRNA-mediated knockdown in COS7 cells, 3 μl Lipofectamine™ RNAiMAX Transfection Reagent was used with the following siRNA: adaptor related protein complex 3 delta 1 subunit (AP3D1) (Dharmacon, ON-TARGETplus SMART siRNA, catalogue no. LQ-016014-00-0002) at a final concentration of 10 nM. Knockdown efficiency was validated by immunoblotting for AP3D1, 48 hours after transfection.

### Neuronal co-culture device

tdTomato-tagged TH DA neurons (day 30) were replated on one side of the two-chamber microfluidic compartmentalization device (OMEGA^4^, eNuvio), where only axonal processes can migrate through the microfluidic channels connected to the adjacent chamber. After an additional 25 days in the co-culture device, frozen iPSC-derived medium spiny neurons (MSN) from Brainxell were plated on the other half of the device (where only the axons of DA neurons are present). The DA-MSN co-cultures were then fixed 7-10 days later for immunofluorescence or EM analysis.

### Immunofluorescence, live imaging and fluorescent microscopy

For all imaging, cells were seeded on glass-bottom matTek dishes (MatTek corporation). For immunofluorescence visualization, cells were fixed with 4% (v/v) paraformaldehyde (Electron Microscopy Sciences) in 1x phosphate-buffered saline (PBS) for 20 mins followed by three washes in PBS. Cells were permeabilized with 0.25-0.5% (v/v) Triton X-100 in PBS for 5 mins followed by three washes in PBS. Cells were then incubated with fresh 1 mg/ml sodium borohydride (Sigma-Aldrich) in PBS for 7 mins to reduce autofluorescence, and then washed thrice in PBS. They were further blocked for 30 min in 5% bovine serum albumin (BSA, Sigma-Aldrich) in PBS and then incubated overnight at 4 °C with the primary antibodies listed in Table S1. Subsequently, cells were washed with PBS thrice the following day and incubated with Alexa Fluor-conjugated secondary antibodies (Thermo Fisher Scientific) for 1 h at room temperature, followed by three washes in PBS. DAPI (Thermo Fisher Scientific) was used for nuclear staining.

For calcium imaging, cells were incubated with FLUO-4 (Thermo Fisher Scientific) at a final concentration of 1 μM for 15 mins followed by 2 washes in neuronal media. Transfections were carried out as described above. For live imaging, cells were maintained in Live Cell Imaging buffer (Life Technologies) for COS7 cells, while both i^3^Neurons and DA neurons were maintained in CM and NB/B27 media, respectively, in a caged incubator with humidified atmosphere (5% CO2) at 37°C. The Yokogawa spinning disk field scanning confocal system with microlensing (CSU-W1 SoRa, Nikon) controlled by NIS elements (Nikon) software was used for neuronal imaging. Excitation wavelengths between 405-640 nm, CFI SR Plan ApoIR 60XC WI objective lens and SoRa lens-switched light path at 1x, 2.8x or 4x were used. SoRa images were deconvolved using the Batch Deconvolution (Nikon) software. The Andor Dragon Fly 200 (Oxford Instruments) inverted microscope equipped with a Zyla cMOS 5.5 camera and controlled by Fusion (Oxford Instruments) software was used for imaging of COS7 cells. Excitation wavelengths between 405-640 nm and Plan Apo 60x (1.45-NA) objective lens were used to visualize COS7 cells.

### Immunoblotting

i^3^Neurons, DA neurons and MSNs were grown on six-well plates (3-5 × 10^5^ cells/well). After differentiation in their respective maturation media, neurons were washed with ice-cold PBS and then lysed in 1xRIPA lysis buffer (10X RIPA lysis buffer, Sigma-Aldrich) supplemented with cOmplete™ EDTA-free protease inhibitor cocktail (Roche) and PhosSTOP phosphatase inhibitor cocktail (Roche), followed by centrifugation at 13,000 × g for 6 min.

The supernatant was collected and incubated at 95 °C for 5 min in SDS sample buffer containing 1% 2-mercaptoethanol (Sigma). The extracted proteins were separated by SDS-PAGE in Mini-PROTEAN TGX precast polyacrylamide gels (Bio-Rad) and transferred to nitrocellulose membranes (Bio-Rad) at 100 V for 1 h or 75 V for 2 h (for high molecular weight proteins: >150 kDa). Subsequently, the nitrocellulose membranes were blocked for 1 h with 5% non-fat milk (AmericanBIO) in TBST (tris-buffered saline [TBS] + 0.1% tween 20), then incubated overnight at 4 °C with primary antibodies and then incubated with IRDye 680RD or 800CW (LI-COR) secondary antibodies (1:8000) for 1 h at room temperature in TBST. For VGLUT2 immunoblotting, blot was re-probed after stripping with Restore™ PLUS Western Blot Stripping Buffer (Thermo Fischer Scientific) for 10 min, followed by three washes with TBST three times, and re-blocked as described above. Finally, blots were imaged using the Gel Doc imaging system (Bio-Rad) using manufacturer’s protocols.

### EM sample preparation and CLEM

COS7 cells were plated on 35 mm gridded, glass-bottom MatTek dish (P35G-1.5-14-CGRD) and transfected as described above. Cells were prefixed with 4% (v/v) PFA in Live Cell Imaging Buffer (Life Technologies) for 15 min followed by three times washing with the same buffer. Regions of interest were selected by fluorescence light microscopy imaging and their coordinates were identified using phase contrast. Cells were further fixed with 2.5% glutaraldehyde in 0.1 M sodium cacodylate buffer for 1 h at room temperature followed by 4 times washing with 0.1M sodium cacodylate buffer for 5 min each. Cells were postfixed in 2% OsO4 and 1.5% K4Fe(CN)6 (Sigma-Aldrich) in 0.1 M sodium cacodylate buffer on ice, then washed 4 times with Milli-Q water for 5 min each. DA neurons were prefixed with 4% PFA as described, further fixed with 2.5% glutaraldehyde and 2mM CaCl2 in 0.15M sodium cacodylate buffer for 1 h at room temperature, and postfixed in 2% OsO4, 1.5% K4Fe(CN)6 and 2mM CaCl2 in 0.15 M sodium cacodylate buffer for 1 h on ice. The specimens were stained with 2% uranyl acetate dissolved in Milli-Q water for 1 h. After washing 4 times with Milli-Q water, specimens were dehydrated in a graded series of ethanol (50%, 75%, and 4 times in 100% for 5 min each), followed by embedding in Embed 812. Regions of interest were relocated based on the pre-recorded coordinates, sectioned and imaged.

For HRP uptake assay, neurons were incubated with 15 μg/ml HRP conjugated CTX (Thermo Fischer Scientific) at 37 °C. HRP reaction was carried out with diaminobenzidene (Sigma-Aldrich, D-5637) (0.5 mg/ml) and H2O2 (JTBaker, 2186-01) (0.01%) in 0.1 M ammonium phosphate buffer (pH7.4) after the glutaraldehyde fixation step. Ultrathin sections (50–60 nm) were observed in Talos L120C TEM microscope at 80 kV. Images were taken with Velox software and a 4k × 4 K Ceta CMOS Camera (Thermo Fischer Scientific). Except when noted, all EM reagents are from Electron Microscope Sciences.

### Fluorescence image analysis, EM and statistics

Images were pseudocolor-coded, adjusted for brightness and contrast, cropped and/or rotated using the open-source image processing software FIJI (ImageJ) (Schindelin et al., 2012). For Figure 1B, semi-automatic segmentation using Cellpose in Napari was used to segment the nucleus (DAPI staining), followed by manual identification of βIII-tubulin and/or TH signals. Immunoblot data were processed using Image Lab software (Bio-Rad) and quantified using the “Gels” plugin in FIJI. The methods for statistical analysis and sizes of the samples (n) are specified in the results section or figure legends for all of the quantitative data. Student’s t-test or Mann–Whitney test was used when comparing two datasets.

Quantification of vesicle diameter was performed using randomly sampled images from clustered vesicles in classical synapses and bouton-like structures for DA neurons, and large droplets in COS7 cells. The diameter of synaptic vesicles was measured from the average of long and short axes. Measurements were performed using FIJI (ImageJ). Differences were accepted as significant for P < 0.05. Prism version 9 (GraphPad Software) was used to plot, analyze and represent the data.

### Statistical analysis

The methods for statistical analysis and sizes of the samples (n) are specified in the results section or figure legends for all quantitative data. Student’s t-test or Mann–Whitney test was used when comparing two datasets. Differences were accepted as significant for P < 0.05. Prism version 9 (GraphPad Software) was used to plot, analyze and represent the data.

### Data availability

All data generated or analyzed during this study are included in this published article (and its Supplementary Information files: Supplementary Material). Raw datasets generated during and/or analyzed during the current study are available from the corresponding authors on request.

### Code availability

Custom-written code used to analyze the data in the current study is available from the corresponding authors on request.

---

**Table S1.**
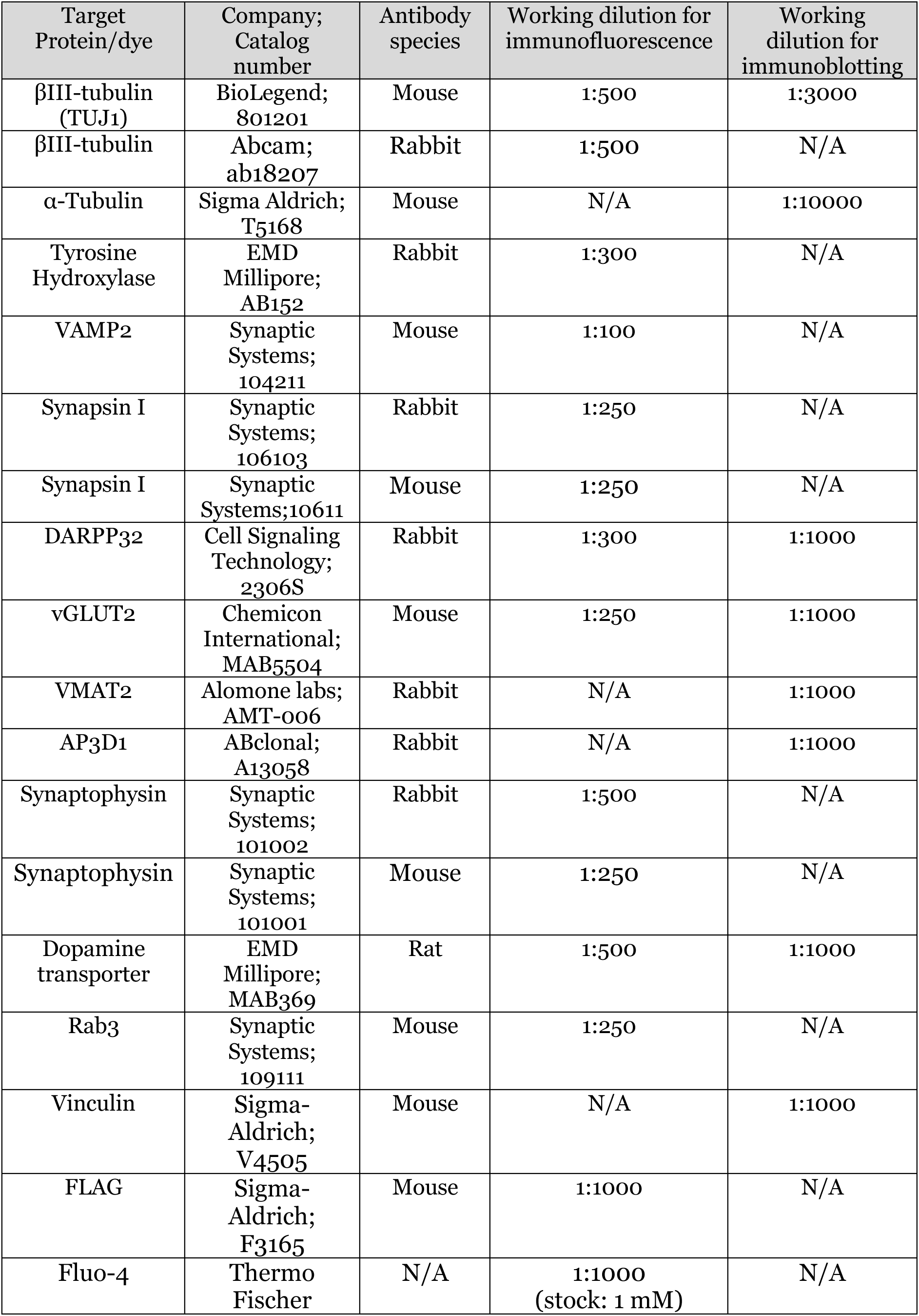
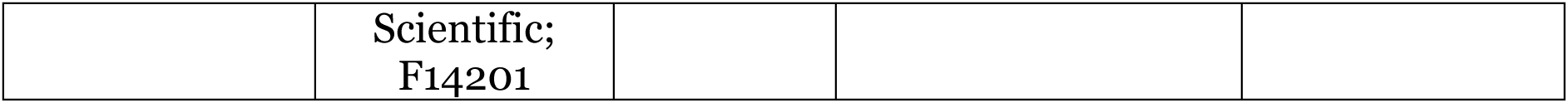
List of antibodies/dyes used in this study.

**Supplementary Figure 1:**
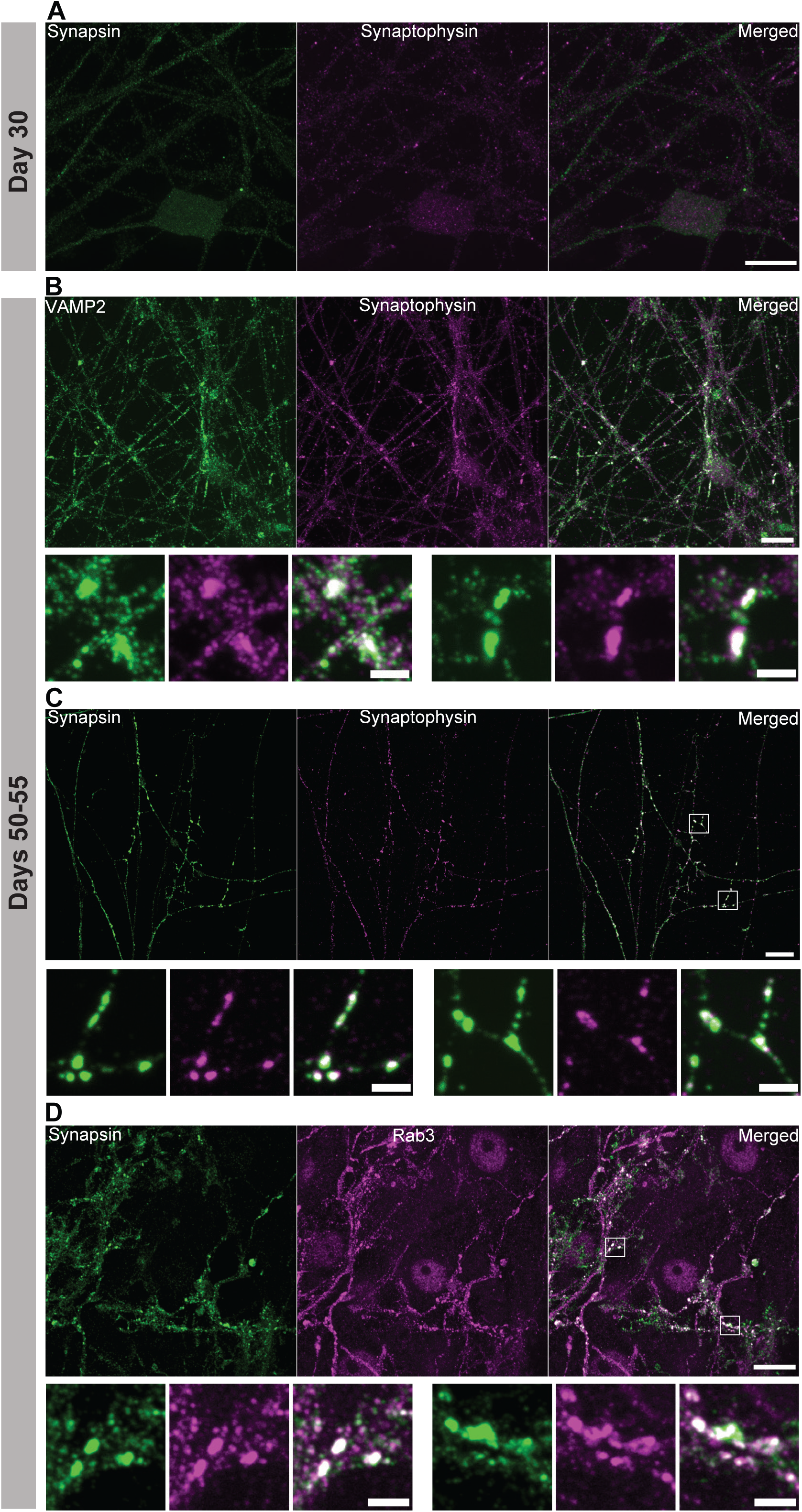
Colocalization of other synaptic markers in DA neurons. (A) Representative fluorescence image of DA neurons (day 30) immunolabeled with antibodies against synapsin (green) and synaptophysin (magenta). (B-D) At day 50-55, DA neurons were immunolabeled with antibodies directed to the following synaptic protein combinations: (B) VAMP2 (green) and synaptophysin (magenta), (C) synapsin (green) and synaptophysin (magenta) or (D) synapsin (green) and Rab3 (magenta). Scale bars, 10 μm. Two high magnification images are shown below each panel showing overlapping fluorescence intensities (white) for the aforementioned synaptic markers. Scale bars, 2 μm.

**Supplementary Figure 2:**
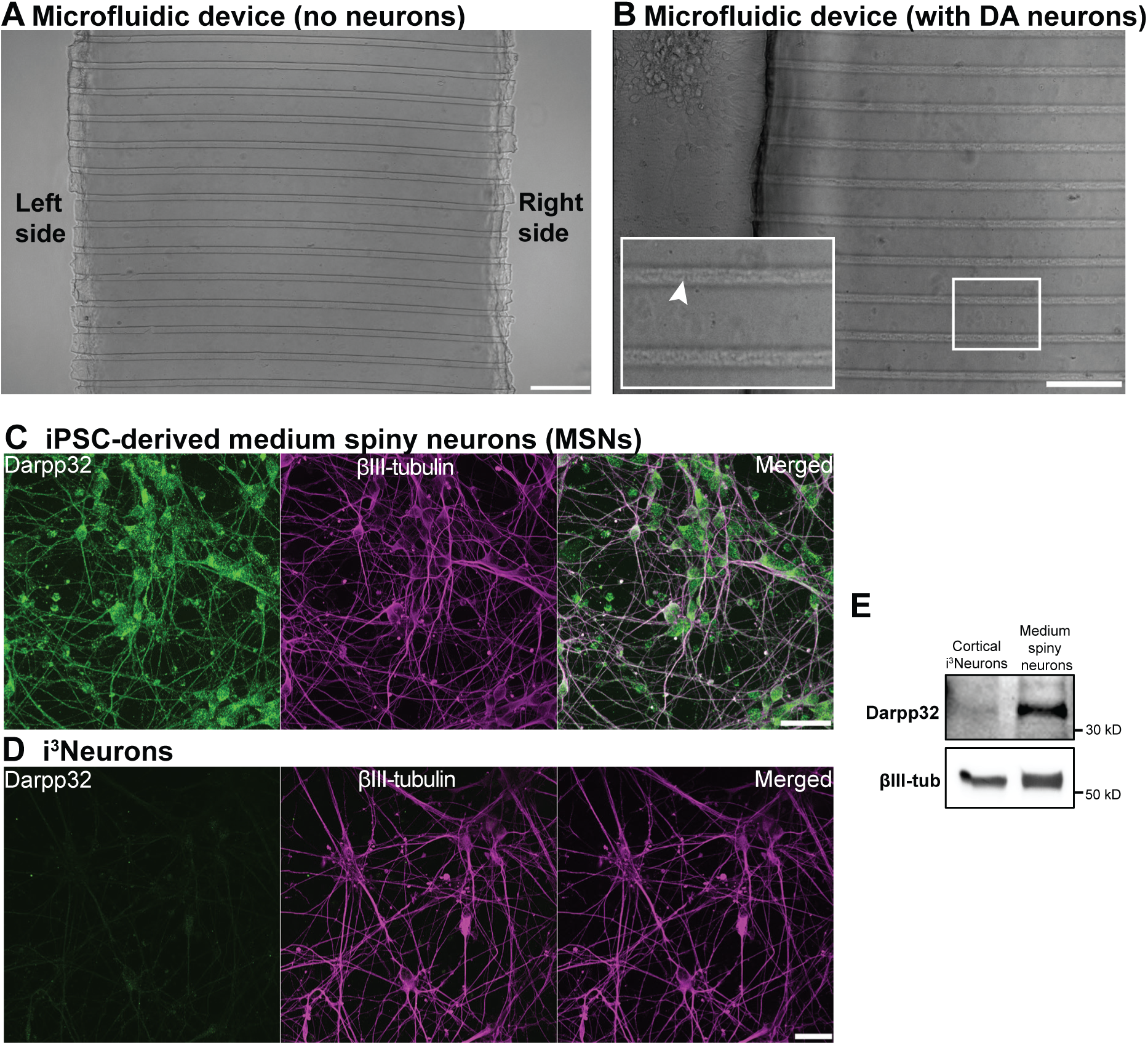
Co-culture microfluidic compartmentalization device set-up. (A and B) A compartmentalization device with two chambers (left and right sides) interconnected by a narrow microfluidic channel visualized before (A) and after seeding of DA neurons on one side of the chamber (B). Inset shows arrows (white) indicating axonal outgrowth in the microchannels. Scale bars, 100 μm. (C and D) iPSC-derived MSNs (day 20, B) from Brainxell and i^3^Neurons (day 19, C) were immunolabeled with antibodies directed against darpp32 (green) and βIII-tubulin (magenta). Note the strong immunoreactivity of darpp32 in MSNs but not in i^3^Neurons. Scale bars, 10 μm. (E) Anti-darpp32 and βIII-tubulin (loading control) western blot of i^3^Neurons and MSNs (20 days in cultures).

**Supplementary Figure 3:**
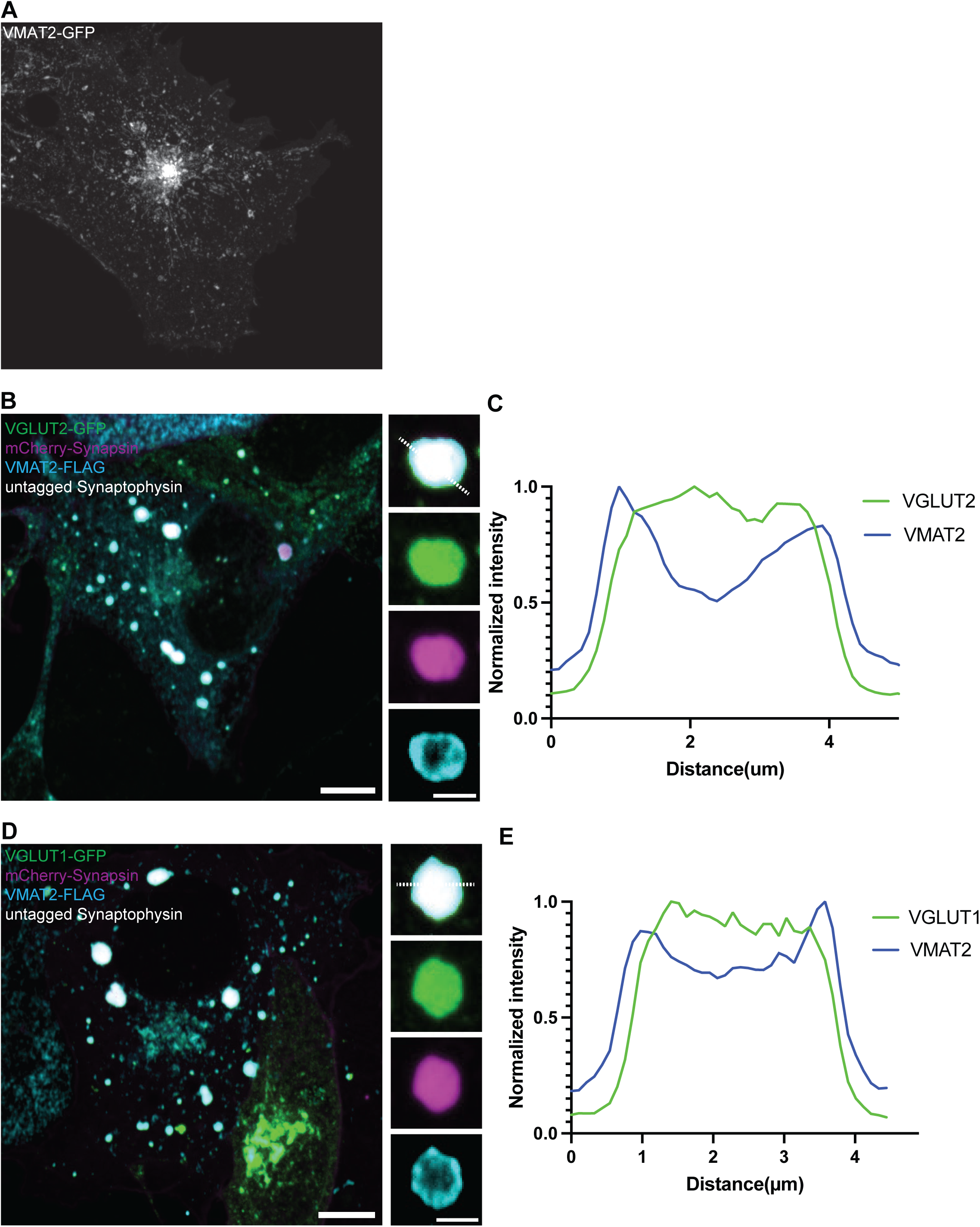
VGLUT1 and VGLUT2 show preferential recruitment to synaptophysin-synapsin over VMAT2-synapsin vesicle clusters. (A) Overexpression of VMAT2-GFP (white) in COS7 cell show Golgi and vesicle puncta stainings. (B) Co-expression of VGLUT2-GFP (green), mCherry-synapsin (magenta), synaptophysin (untagged) and VMAT2-FLAG (cyan) in COS7 cells show preference of VGLUT2-GFP fluorescence signals in the synaptophysin-synapsin vesicle clusters. Scale bars, 10 μm; inset: 2 μm. (C) The corresponding line-scan analysis from the inset in B (dotted white line) is shown for both VMAT2-FLAG and VGLUT2-GFP. (C) Co-expression of VGLUT1-GFP (green), mCherry-synapsin (magenta), synaptophysin (untagged) and VMAT2-FLAG (cyan) in COS7 cells show preference of VGLUT1-GFP fluorescence signals in synaptophysin-synapsin vesicle clusters. Scale bar, 10 μm; inset: 2 μm. (E) The corresponding line-scan analysis from the inset in D (dotted white line) is shown for both VMAT2-FLAG and VGLUT1-GFP. VMAT2-FLAG was revealed by immunofluorescence of antibody against FLAG.

**Supplementary Figure 4:**
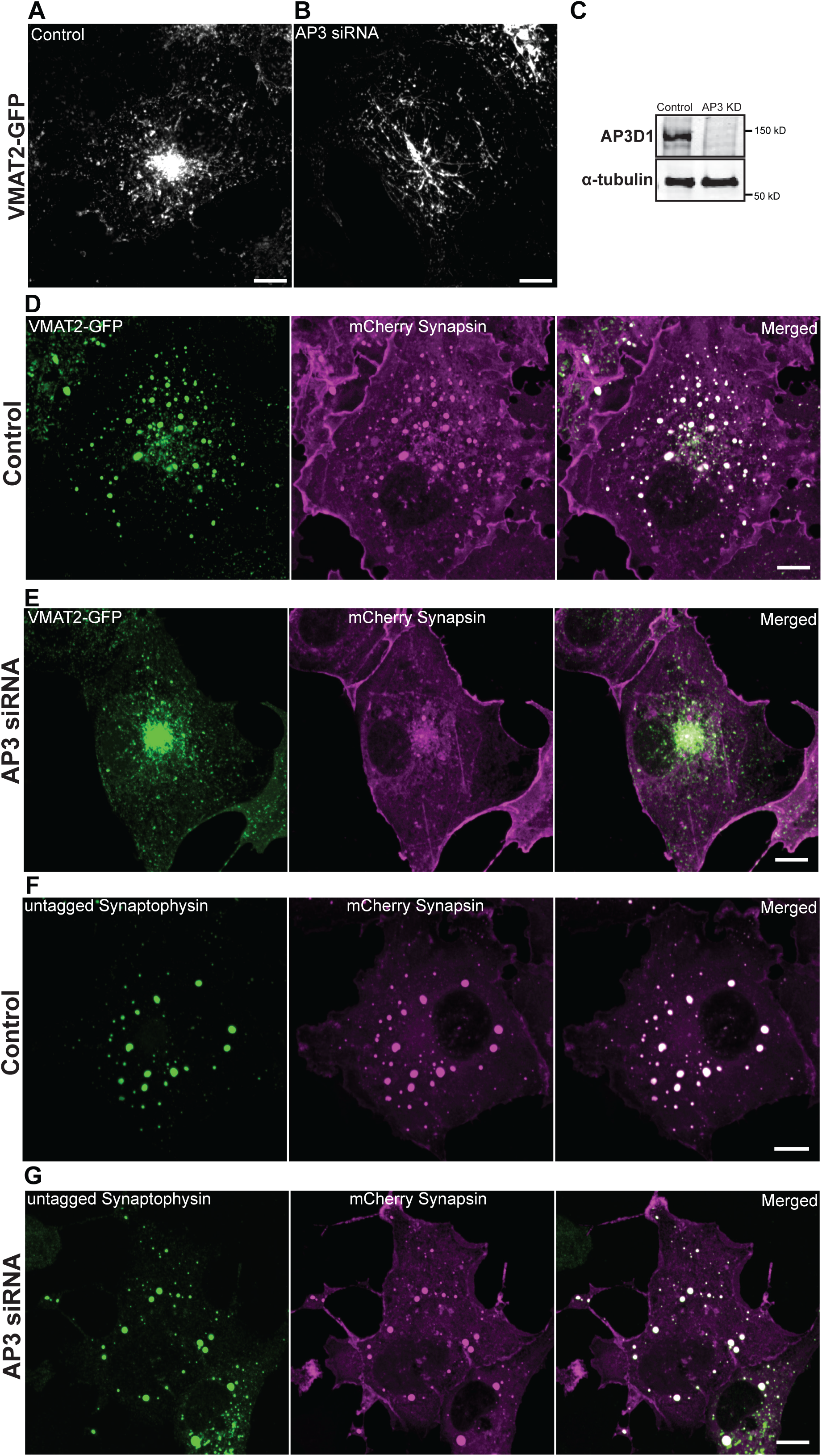
AP3 knockdown abolished VMAT2-synapsin but not synaptophysin-synapsin vesicle clusters. (A and B) Representative fluorescence images of COS7 cells transfected with VMAT2-GFP and either control siRNA (A) or AP3D1 siRNA (B) visualized 48 hours post-transfection. Scale bars, 10 μm. (C) Anti-AP3D1 and α-tubulin (loading control) western blot of control and AP3 KD siRNA-treated COS7 cells 48 hours post-transfection. (D and E) COS7 cells transfected with VMAT2-GFP (green) and mCherry-synapsin (magenta) treated with control siRNA (D) or AP3D1 siRNA (E). (F and G) COS7 cells transfected with untagged synaptophysin (green) and mCherry-synapsin (magenta) treated with control siRNA (F) or AP3D1 siRNA (G). Untagged synaptophysin was revealed by immunofluorescence. Images from D-G were taken 48 hours after transfection. Note the lack of VMAT2-synapsin droplets in E. Scale bars, 10 μm.

